# Population structure and genetic connectivity of the scalloped hammerhead shark (*Sphyrna lewini*) across nursery grounds from the Eastern Tropical Pacific: implications for management and conservation

**DOI:** 10.1101/2022.02.22.481487

**Authors:** Mariana Elizondo-Sancho, Yehudi Rodríguez-Arriatti, Federico J. Albertazzi, Adrián Bonilla-Salazar, Daniel Arauz, Randall Arauz, Elisa Areano, Cristopher G. Avalos-Castillo, Óscar Brenes, Elpis J. Chávez, Arturo Dominici-Arosemena, Mario Espinoza, Maike Heidemeyer, Rafael Tavares, Sebastián Hernández

## Abstract

Defining demographically independent units and understanding gene flow between them is essential for managing and conserving exploited populations. The scalloped hammerhead shark, *Sphyrna lewini*, is a coastal semi-oceanic species found worldwide in tropical and subtropical waters. Pregnant females give birth in shallow coastal estuarine habitats that serve as nursery grounds for neonates and small juveniles, and adults move offshore and become highly migratory. We evaluated the population structure and connectivity of *S. lewini* in coastal areas across the Eastern Tropical Pacific (ETP) using both sequences of the mitochondrial DNA control region (mtCR) and nuclear-encoded microsatellite loci. The mtCR defined two genetically discrete geographic groups: the Mexican Pacific and the central-southern Eastern Tropical Pacific (Guatemala, Costa Rica, Panamá, and Colombia). Overall, the mtCR data showed low levels of haplotype diversity ranged from 0.000 to 0.608, while nucleotide diversity ranged from 0.000 to 0.0015. A more fine-grade population structure analysis was detected using microsatellite loci where Guatemala, Costa Rica, and Panamá differed significantly. Genetic diversity analysis with nuclear markers revealed an observed heterozygosity ranging from 0.68 to 0.71 and an allelic richness from 5.89 to 7.00. Relatedness analysis revealed that individuals within nursery areas were more closely related than expected by chance, suggesting that *S. lewini* may exhibit reproductive philopatric behaviour within the ETP. Findings of at least two different management units, and evidence of philopatric behaviour call for intensive conservation actions for this critically endangered species in the ETP.

## Introduction

Delimiting demographically independent populations and understanding their level of genetic diversity and connectivity is central to managing and conserving endangered and exploited species (1–3). In aquatic ecosystems, animals that occupy high trophic positions generally exhibit high extinction risks due to their large size, life-history characteristics, and the exploitation rates they are subjected to (4,5). Sharks are one of the most threatened groups of marine fishes globally, mainly due to overfishing and habitat degradation which has increased dramatically over the past 20 years (4,6,7). The effects of population-level declines are of major concern in conservation since small populations suffer from inbreeding and genetic drift, which may lead to loss of genetic diversity and compromise the ability of a population to evolve and cope with environmental change (8).

The scalloped hammerhead shark *Sphyrna lewini* (Griffith and Smith, 1834), is a large (up to 420 cm total length, TL), viviparous, coastal semi-oceanic species found worldwide in tropical and sub-tropical waters (9). As occurs with many other shark species, *S. lewini*, has low resilience to overfishing due to its slow growth, late sexual maturity, and long gestation periods (10–12). Throughout its distribution, *S. lewini* has experienced severe population declines (7,13–16), leading to its listing as Critically Endangered by the International Union for the Conservation of Nature (IUCN) Red List (17). The scalloped hammerhead shark has a complex life history in which pregnant females give birth in shallow coastal estuarine habitats that serve as nursery grounds for neonates and small juveniles during their first years of life (18,19). Eventually, large juveniles and adults move offshore and become highly migratory, often schooling around seamounts and near continental shelves(20,21).

The dichotomy between breeding in coastal reproductive habitats and the long-range dispersal of adults displayed by shark species such as *S.lewini* may result in complex population structure (22). Six distinct population segments of *S. lewini* have been distinguished globally, defined within 1) the North West Atlantic and Gulf of Mexico, 2) Central and South West Atlantic, 3) Eastern Atlantic, 4) Indo-West Pacific, 5) Central Pacific, and 6) in the Eastern Pacific (16,23). Throughout the Eastern Tropical Pacific (ETP) Ocean, *S. lewini* is subject to high fishing pressures, in both coastal and oceanic waters. Neonates and juveniles are susceptible to shrimp trawl and small-scale fisheries inshore, whereas adults are a frequent by-catch in pelagic longline and purse-seine fisheries that operate near seamounts and oceanic islands (16,24,25).

Genetically discrete groups are created by reproductive behaviors that segregate populations, which cause allelic frequencies to diverge in time (26). Natal philopatry is described as a reproductive behavior in which organisms return to their birthplace to reproduce or give birth (27). This behavior has been observed in several species of sharks, including the great white shark (*Carcharodon carcharias*), ma ko shark (*Isurus oxyrinchus*), lemon shark (*Negaprion brevirostris*), blacktip shark (*Carcharhinus limbatus*), and sand tiger shark (*Charcharhinus taurus*) (28–33). For a species that uses coastal habitats as nursery areas, such as *S. lewini*, natal philopatry could contribute to the development of genetically discrete groups, where intrinsic reproduction and recruitment may result in population structure at smaller geographic scales than would be expected based on the mobility of the organism (34,35).

To date, studies investigating the genetic structure of *S. lewini* in the ETP have been either been limited to small geographic areas (7,36) or they used a relatively small sample size (37). Given the limited data on the population structure of *S. lewini* and the high fishing pressure that this species is currently under throughout the ETP, sorting out the genetic diversity to assess population structure and connectivity in potential nursery areas of the region, is crucial to develop effective management and conservation strategies. The information pertaining to 14 microsatellite loci (bi-parentally inherited) and a 489-bp fragment of the mitochondrial DNA control region (maternally inherited) was used to analyze the genetic variation and differentiation in samples sourced from young of the year (YOY) *S. lewini* individuals on coastal sites in the ETP. This study (i) assessed the genetic diversity of *S. lewini* in coastal sites of the ETP, (ii) determined the population structure of *S. lewini* within the ETP, and (iii) evaluated the potential role of natal philopatry in the population dynamics of *S. lewini* within the ETP.

## Materials and methods

### Study region

The study region comprises the majority of the ETP (Fig. 1), from the coast of Central America and South America to 140°W (38). The ETP includes a complex diversity of coastal environments and oceanic islands with oceanographic conditions that vary seasonally, annually and over longer time scales (39). Coastal sampling sites were comprised of estuarine systems with predominant mangrove vegetation and muddy coasts 1) Las Lisas and Sipacatein Guatemala (GUA); 2) Coyote (COY) and Ojochal (OJO/COS) in Costa Rica; 3) and Punta Chame in Panamá (PAN) (Fig. 1). Previously collected molecular data from coastal areas in México and Colombia were included in the analysis to cover a broader geographic range. Mexican sites included: Nayarit (NAY), Oaxaca (OAX), Michoacan (MCH), Baja California (BJC), Chiapas (CHP), and Sinaloa (37); Colombia sites included: Port Buenaventura (PTB), Utria (UTR), Sanquianga (SNQ) and Malpelo Island (MLP) (36) (Fig. 1).

**Figure 1.**
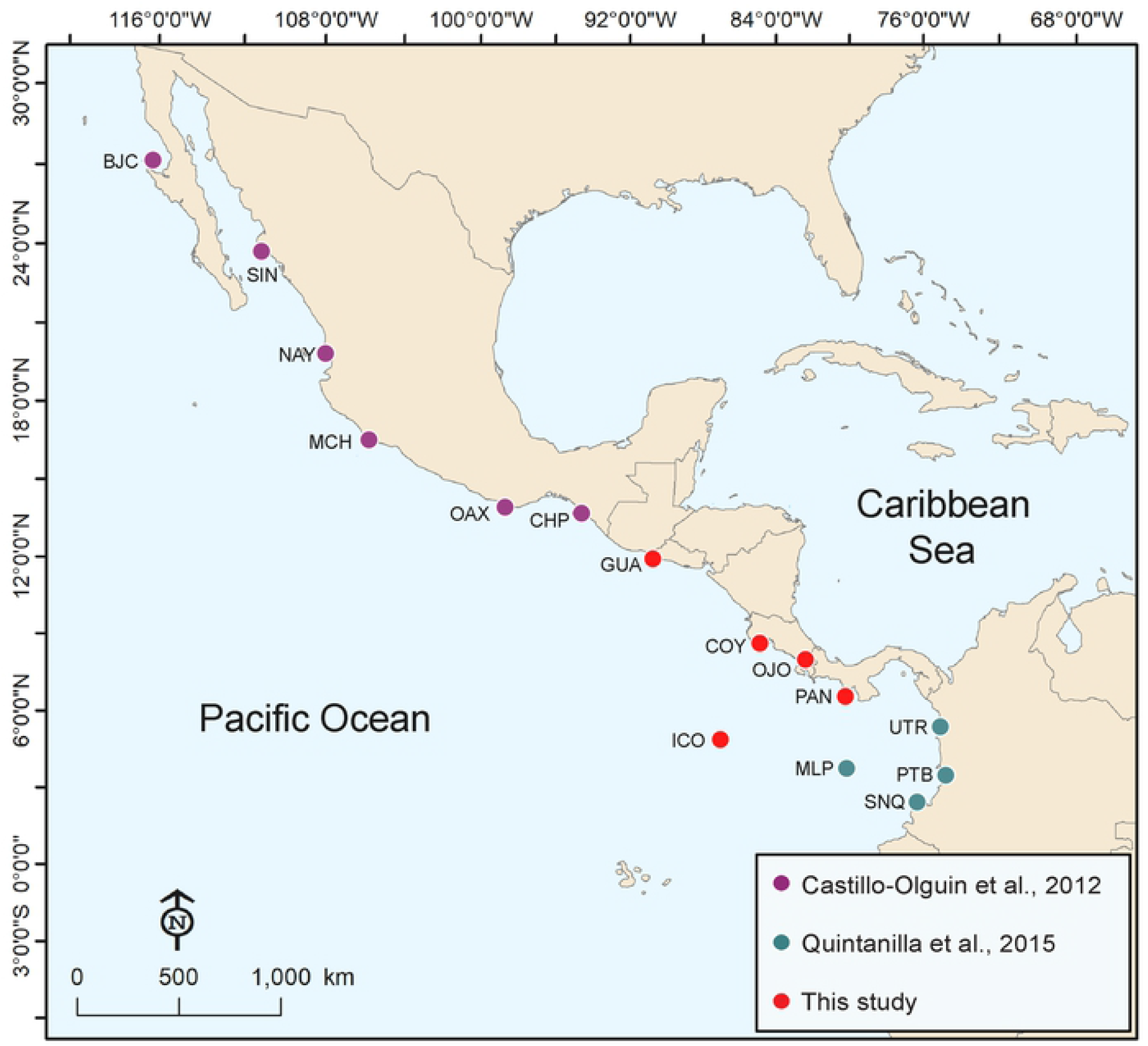
Location of sampling sites of *Sphyrna lewini* individuals in the Eastern Tropical Pacific. Sampling sites: Guatemala (GUA), Ojochal (OJO), Coyote (COY), Cocos Island (ICO), Panama (PAN), Nayarit (NAY), Oaxaca (OAX), Michoacan (MCH), Baja California (BJC), Chiapas (CHP), Sinaloa (SIN), Port Buenaventura (PTB), Sanquianga (SNQ), Utria (UTR), Malpelo Island (MLP).

### Sample collection

Tissue samples of juvenile *S. lewini* (30-50 cm TL) were collected from artisanal fisheries operating along the coast in Costa Rica (N = 78), Panamá (N = 65) and Guatemala (N = 72) throughout 2017 and 2018. In addition, samples from adults (63-108 cm TL) were collected opportunistically in Cocos Island (N = 16) with a biopsy dart during scientific cruises conducted in 2008. The use of tissue samples for this study was reviewed by the National Commission for the Management of Biodiversity (CONAGEBIO) of Costa Rica. The technical office of CONAGEBIO emitted the research permit R-CM-VERITAS-001-2021-OT-CONAGEBIO. The Ministry of Environment of Panamá issued the research permits SEX/A-61-19 and SEX/A-108-17 and the National Council of Protected Areas of Guatemala issued the research license no. I-DRSO-001-2018. Fin and muscle tissue was preserved in 95% ethanol and stored at −20° C. Total DNA was extracted from 25 mg of tissue using the phenol-chloroform protocol (40) and with Promega’s Wizard^®^ Genomic DNA Purification Kit.

### Amplification and sequencing of mitochondrial DNA

The mitochondrial DNA control region (mtCR) was amplified and sequenced for a total of 231 *S. lewini* individuals using species specific primers designed in Geneious Pro v6.0.6 Bioinformatics Software for Sequence Data Analysis (41). Forward (3’ AAGGGTCAACTTCTGCCCT 5’) and reverse (3’AGCATGGCACTGAAGATGCT 5’) primers were designed based on the whole mitochondrial genome of *S. lewini* deposited in Genbank (Accession number: JX827259). PCR amplification was conducted using a Veriti™ Thermal Block (Applied Biosystems, USA) with a total volume of 15μL containing 67 mM Tris-HCl pH 8.8, 16mM (NH_4_)_2_SO_4_, 2.0 mM MgCl_2_, 20 mM dNTPs, 10 μM of each primer, 0.4 units of Dream Taq DNA Polymerase (5U/ μl), and 1 μl of DNA (20-40 ng/μl). The PCR thermal profile included initial 5 min denaturation at 94°C, 30 cycles of 30 s at 94°C, 30 s at 59°C and 1.5 min at 72°C, followed by a final extension for 10 min at 72°C. PCR products and the corresponding negative control were visualized in UV light after electrophoresis in 1.2% agarose gel. PCR products were purified and sequenced in both directions using an ABI 3100 automated sequencer.

### Amplification and genotyping of microsatellite loci

A total of 169 samples of *S. lewini* were genotyped for 14 microsatellite loci previously described by Nance et al. (2009 *lewini* (Guatemala = 52; Costa Rica = 50; Panamá =5; Cocos Island =16). Forward primers were marked with an M13 tail (5’-TGT AAA ACG ACG GCC AGT-3’) (42). Microsatellite amplification was conducted using a nested PCR in a total volume of 15 μL with 1-2 μL of DNA (10-30 ng), 0.1 μM forward primer, 0.4 μM primer reverse, 0.4 μM M13 primer (6FAM, VIC or NED), 0.2 mM dNTPs, 2 mM MgCl_2_, 0.04 units of Dream Taq DNA Polymerase (5U/μL), 1X Buffer and water. PCR conditions consisted of an initial 2 min denaturalization at 94°C, followed by 32 cycles of 30 s at 94°C, 30s of 57°C (Sle25, Sle77), 59°C (Sle45, Sle59, Sle33, Sle53), 60°C (Sle54, Sle13, Sle18, Sle27, Sle81, Sle71, Sle86, Sle38) 1 min a 72°C, followed by 8-20 cycles of 30 s at 94 °C, 30 at 53°C, 30 at 72°C and a final extension of 2 min at 72 °C. PCR products and corresponding negative control were verified by electrophoresis in 1.2% agarose gel and visualized using UV light. Fragment size analysis was done using an ABI 3100 automated sequencer.

### Mitochondrial DNA analysis

Collected in this study from coastal sites of the ETP and Cocos Island, 229 sequences from the mtCR of *S.lewini* were analyzed. An alignment of 489bp was carried out with the MUSCLE algorithm on GeneiousPro Bioinformatics Software for Sequence Data Analysis (41). For a broader geographic range, 205 additional sequences previously published were retrieved from GenBank^®^ (Table S1) and added to the alignment from the coast of Colombia (36) and México (37). Arlequin 3.5 Software (43) was used to calculate the number of haplotypes (H), polymorphic sites (S), nucleotide diversity (π), haplotype diversity (hd) and nucleotide base composition. To examine relationships among haplotypes a haplotype network was drawn in Haploview Bioinformatics software (44) which was based on phylogenetic reconstructions carried out for maximum likelihood in RAxML-HPC2 8.2.12 on XSEDE (45) (available at https://www.phylo.org) (46). The maximum likelihood analysis was carried out using GTR+ Gamma and 1000 Bootstrap iterations.

Arlequin 3.5 software (43) was used to estimate the Population’s genetic structure among geographic areas using Wright’s pairwise fixation index (F_ST_) (47) (10,000 permutations, α = 0.05), with Holm-Bonferroni correction for probabilities (48). A global hierarchical Analysis of Molecular Variance (AMOVA) (Excoffier et al., 1992) was performed to determine the genetic diversity among and within populations using Arlequin 3.5 (10,000 permutations and α = 0.05) (43). An AMOVA was conducted by grouping haplotypes into two groups, one with samples from the Northern ETP and the other with the samples from the Central-southern ETP. A Mantel test (49) was conducted to test the hypothesis that genetic differentiation is due to isolation-by-distance; *Adegenet R* package (50) in R v.4.0.2.(51) was used to evaluate the correlation between Nei’s genetic distance and a matrix of Euclidian geographic distance.

### Microsatellite DNA data analysis

The fragment size of 14 microsatellite loci for each sample was determined by identifying the peaks with GeneMarker^®^ Software 2.6.3. The presence of genotyping errors and null alleles, as well as the frequency of null alleles per locus (r) was evaluated using MICRO-CHECKER v.2.2.3 (52). Deviations from Hardy-Weinberg equilibrium (HW) and linkage disequilibrium (LD) were calculated for each locus and sampling site using GENEPOP v. 4.0 (53) utilizing 10,000 steps of dememorization, 1,000 batches and 10,000 iterations per batch. All probability values were adjusted using the Holm-Bonferroni correction (48). *Adegenet R* package (50) in R v.4.0.2 (51) was used to calculate the number of alleles (A), allelic richness (Ar), expected heterozygosity (HE), observed heterozygosity (HO) and inbreeding coefficient (FIS). A relatedness analysis with a 99% confidence interval was conducted using M-L relate (54) to identify related pairs (full siblings, FS), that were then excluded from further analysis to avoid biased estimations of genetic diversity and population structure.

The package *Related* (55) in R v.4.0.2. (51) was used to conduct the genetic relatedness analysis based on the allele frequencies among all pairs of *S. lewini* individuals within and among sampling sites. The function “comparestimators” was used to select the best relatedness estimator and evaluate the performance of four genetic relatedness estimators. (56–59). This function simulates individuals of known relatedness based on the observed allele frequencies and compares the correlation between observed and expected genetic relatedness for each estimator. The relatedness estimator with the highest correlation coefficient was chosen. To re-check for the possibility of occurrence of related individuals that may bias estimates of genetic diversity and differentiation, the distribution of observed pairwise relatedness values across all individuals was also compared to the values expected between parent-offspring (PO), full-siblings (FS), half-siblings (HS) and unrelated pairs (U). Subsequently, to determine if individuals within sampling sites were more closely related than expected by chance, observed values of relatedness for each sampling site were compared from random mating expectations with the function “grouprel”. This function calculates the average pairwise relationship within each predefined group (i.e., sampling site) as well as an overall within-group relatedness. The expected distribution of average within group relatedness is generated by randomly shuffling individuals using 1,000 Monte Carlo simulations, keeping group size constant. The observed mean relatedness is then compared to the distribution of simulated values to test the null hypothesis of groups being randomly associated in terms of relatedness.

To test for population structure between sample collection areas, pairwise population comparisons of D_EST_ values (60) were obtained using the *DEMEtics* package (61) in R v.4.0.2. Wright’s pairwise fixation index (F_ST_) (47) (20 000 permutations, α = 0.05), with Holm-Bonferroni correction for probabilities (48) was also obtained using Arlequin 3.5 (Excofier and Lischer, 2010). Additionally, the software STRUCTURE v.2.3.4 (Pritchard et al., 2000) was used to identify the clustering of groups of individuals and the admixture with a Markov Chain of Monte Carlo (MCMC) (length burn-in period: 200,000; MCMC: 40,000; 10 K, 10 iterations each). To complement previous population structure analysis, a multivariate approach, Discriminant Analysis of Principal Components (DAPC) (62) was used to identify discrete populations based on geographic region, using the *Adegenet* package in R v.4.0.2. The DAPC summarizes initial genetic data into uncorrelated groups using principal components, then uses discriminant analysis to maximize the among-population variation (63). Cluster assignments were pre-defined corresponding with defined collection locations.

A Mantel test was performed to test the hypothesis of genetic differentiation due to isolation-by-distance; the correlation between Nei’s genetic distance and a matrix of Euclidian geographic distances were evaluated using the *Adegenet* package (50) in R v.4.0.2. Gene flow was analyzed with the “divMigrate” function (64) of the package *diveRsity* (65) in R v. 4.0.2 (51). The “divMigrate” function was used to plot the relative migration levels and detect asymmetries in gene flow patterns, between pairs of population samples (64).

## Results

### Mitochondrial DNA

The nucleotide alignment (434 sequences and 489pb) of mtCR sequences from individuals across the ETP, had a nucleotide base composition of 31.7% A, 24.4% C, 7.8% G, 36.1%T, 16 haplotypes, and 23 polymorphic sites. Sequences from this study are deposited in Genbank, accession numbers: OL692109 - OL692337. There was variation of the genetic diversity of *S. lewini* samples throughout the ETP (Table 1). The haplotype diversity (hd) ranged from 0.000 to 0.608, while nucleotide diversity (π) ranged from 0.000 to 0.0015. The highest genetic diversity was observed in Guatemala (hd = 0.608; π = 0.00015), followed by Malpelo Island (hd = 0.581; π = 0.0012). The lowest genetic diversity was detected in Baja California, Chiapas, and Oaxaca (h = 0.000, π = 0.000). The overall genetic diversity in the ETP was relatively low (hd= 0.3912, π=0.0016).

**Table 1.**
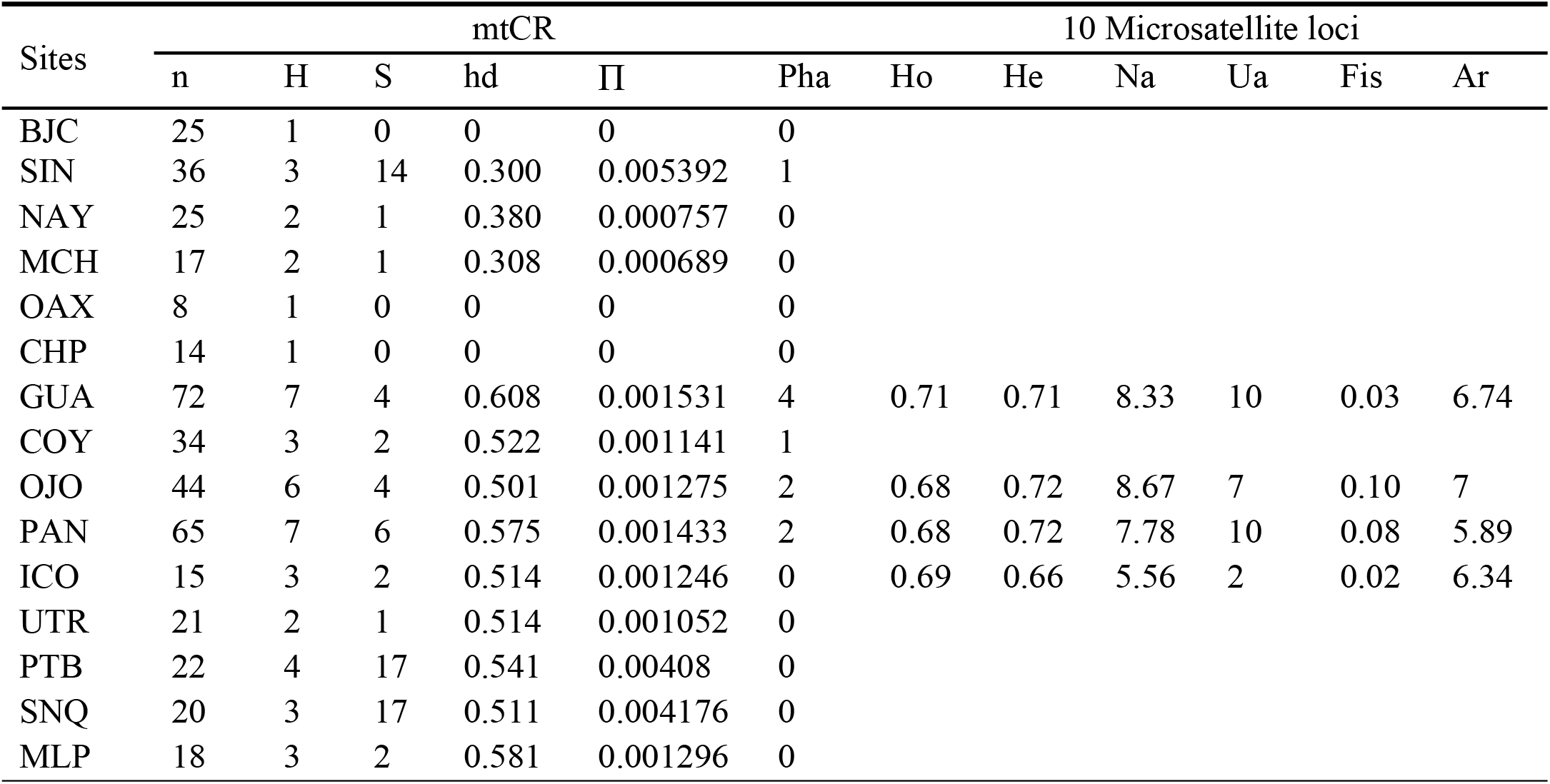
Genetic diversity indexes by sampling site for the mitochondrial control region and 10 microsatellite loci for *Sphyrna lewini* individuals in the Eastern Tropical Pacific. N: number, H: number of haplotypes, S: polymorphic sites, hd: haplotype diversity, π: nucleotide diversity, Pha: number of private haplotypes, Ho: observed heterozygosity, He: expected heterozygosity, Na: number of alleles, Ua: unique alleles, Fis: inbreeding coefficient, Ar: allelic richness. Sampling sites: Baja California (BJC), Sinaloa (SIN), Nayarit (NAY), Michoacan (MCH), Oaxaca (OAX), Chiapas (CHP), Guatemala (GUA), Coyote (COY), Ojochal (OJO), Panama (PAN), Cocos Island (ICO), Utria (UTR), Port Buenaventura (PTB), Sanquianga (SNQ), Malpelo Island(MLP).

A total of 16 haplotypes were found in all samples across the ETP (Fig 2). Thirteen of these haplotypes were sampled out of two or more individuals where Hap5 was the most common haplotype across all sampling sites and detected in 50.4% of all individuals analyzed (Fig. 2, Table S3). Although Hap5 and Hap4 were found in all sampling sites, Hap5 was more common in Mexican sampling sites, while Hap4 appeared to be more common in Central America and Colombia. Ten private haplotypes were detected: GUA (4), OJO (2), COY (1), PAN (2), SIN (1).

**Figure 2.**
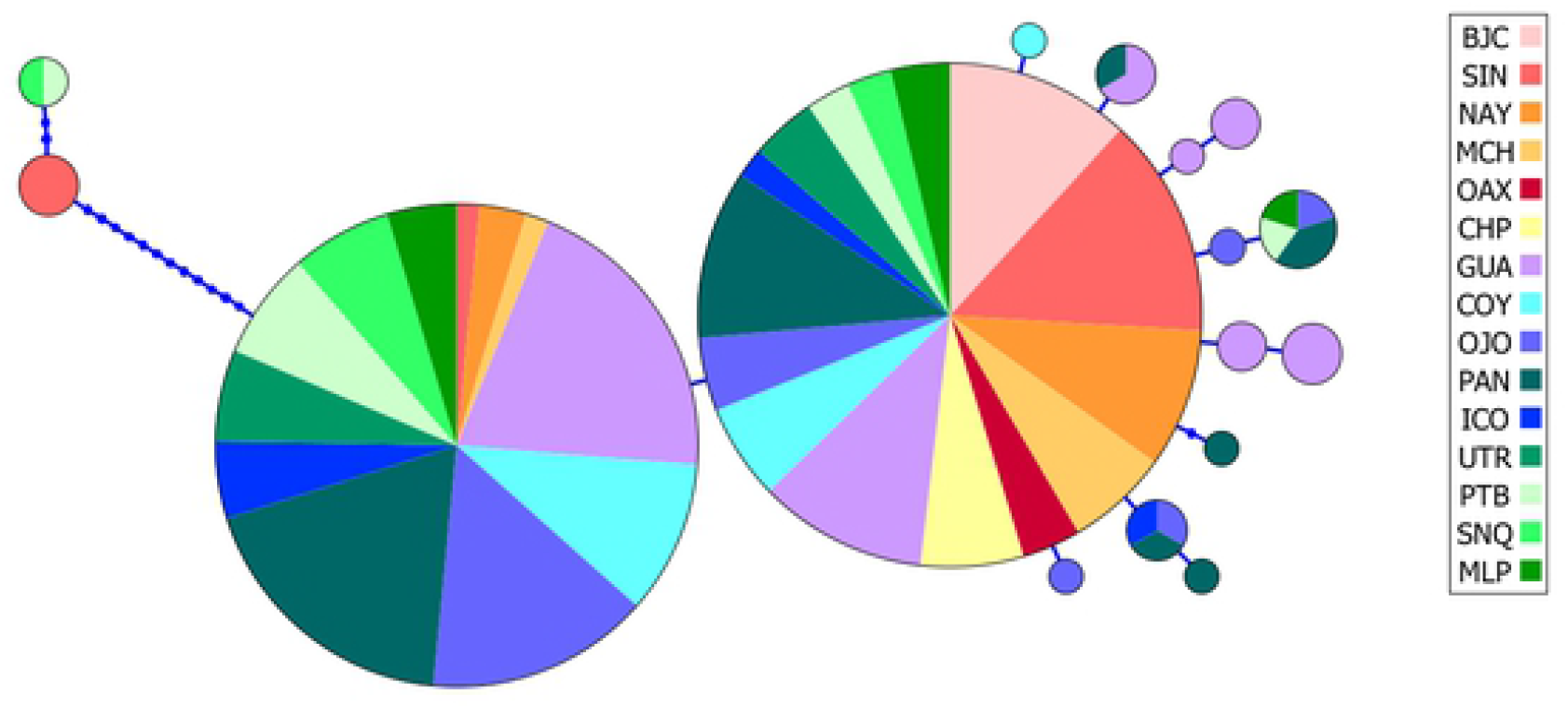
Haplotype network based on mitochondrial control region sequences for *Sphyrna lewini*. Each circle represents a unique haplotype, the colors represent the proportion of individuals from various sampling sites sharing the haplotype. Ticks on connecting lines indicate mutational steps between haplotypes. Sampling sites: Guatemala (GUA), Ojochal (OJO), Coyote (COY), Cocos Island (ICO), Panama (PAN), Nayarit (NAY), Oaxaca (OAX), Michoacan (MCH), Baja California (BJC), Chiapas (CHP), Sinaloa (SIN), Port Buenaventura (PTB), Sanquianga (SNQ), Utria (UTR), Malpelo Island (MLP).

After pairwise Ɵ_ST_ values showing significant genetic differentiation between Northern ETP sampling sites (NAY, OAX, MCH, BJC, CHP, SIN) and Central-southern ETP (GUA, OJO, COY, ICO, PAN, PTB, UTR, SNQ and MLP) (Table 2), two groups were considered for the hierarchal AMOVA: samples from the Northern ETP and samples from Central-southern ETP. Significant levels of population subdivision were found between these two groups, representing 23.7% of the variation found in the mtCR (Table 3). The mtCR Mantel test revealed a significant pattern of isolation-by-distance (r = 0.47, p = 0.002), showing that genetic distance was correlated with geographic distance.

**Table 2.**
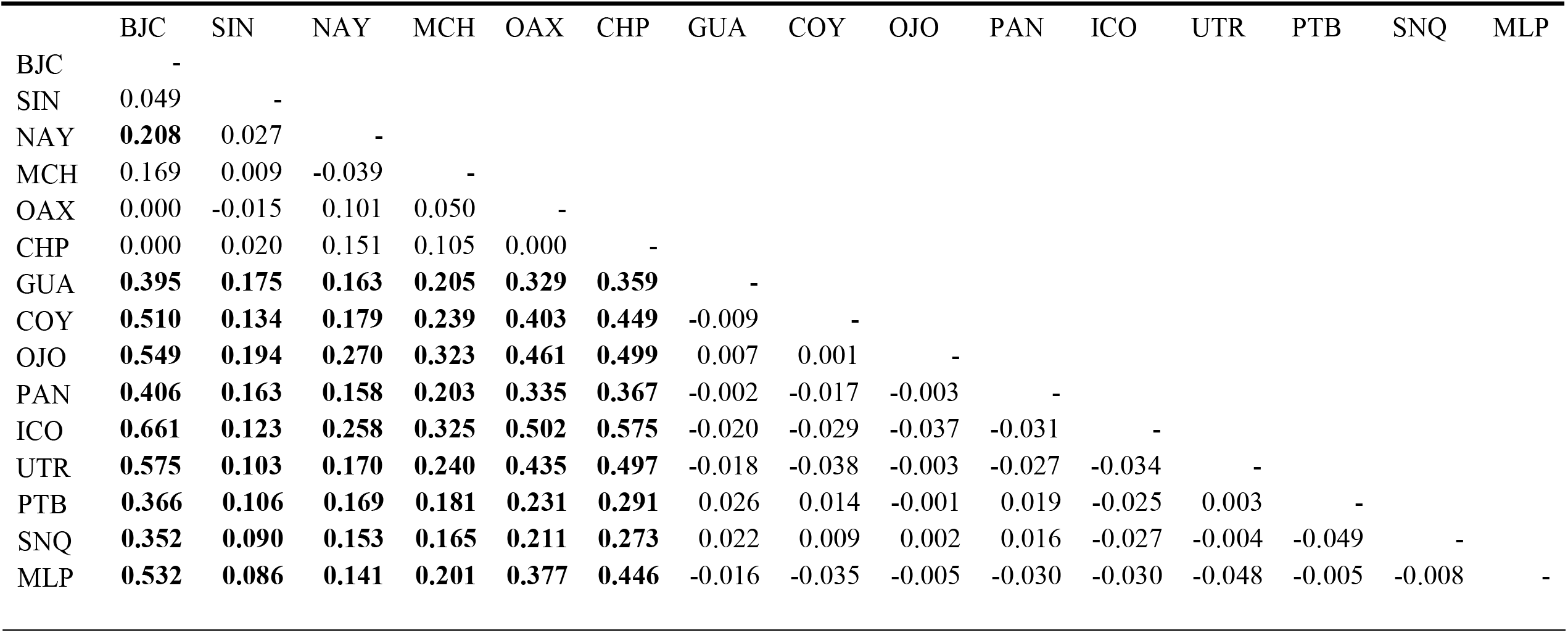
Pairwise F_ST_ of the mitochondrial control region for *Sphyrna lewini* individuals in the Eastern Tropical Pacific. Sampling sites: Sampling sites: Baja California (BJC), Sinaloa (SIN), Nayarit (NAY), Michoacan (MCH), Oaxaca (OAX), Chiapas (CHP), Guatemala (GUA), Coyote (COY), Ojochal (OJO), Panama (PAN), Cocos Island (ICO), Utria (UTR), Port Buenaventura (PTB), Sanquianga (SNQ), Malpelo Island (MLP). Significant values are found in Bold letters.

|  | BJC | SIN | NAY | MCH | OAX | CHP | GUA | COY | OJO | PAN | ICO | UTR | PTB | SNQ | MLP |
| --- | --- | --- | --- | --- | --- | --- | --- | --- | --- | --- | --- | --- | --- | --- | --- |
| BJC | - |  |  |  |  |  |  |  |  |  |  |  |  |  |  |
| SIN | 0.049 | - |  |  |  |  |  |  |  |  |  |  |  |  |  |
| NAY | **0.208** | 0.027 | - |  |  |  |  |  |  |  |  |  |  |  |  |
| MCH | 0.169 | 0.009 | -0.039 | - |  |  |  |  |  |  |  |  |  |  |  |
| OAX | 0.000 | -0.015 | 0.101 | 0.050 | - |  |  |  |  |  |  |  |  |  |  |
| CHP | 0.000 | 0.020 | 0.151 | 0.105 | 0.000 | - |  |  |  |  |  |  |  |  |  |
| GUA | **0.395** | **0.175** | **0.163** | **0.205** | **0.329** | **0.359** | - |  |  |  |  |  |  |  |  |
| COY | **0.510** | **0.134** | **0.179** | **0.239** | **0.403** | **0.449** | -0.009 | - |  |  |  |  |  |  |  |
| OJO | **0.549** | **0.194** | **0.270** | **0.323** | **0.461** | **0.499** | 0.007 | 0.001 | - |  |  |  |  |  |  |
| PAN | **0.406** | **0.163** | **0.158** | **0.203** | **0.335** | **0.367** | -0.002 | -0.017 | -0.003 | - |  |  |  |  |  |
| ICO | **0.661** | **0.123** | **0.258** | **0.325** | **0.502** | **0.575** | -0.020 | -0.029 | -0.037 | -0.031 | - |  |  |  |  |
| UTR | **0.575** | **0.103** | **0.170** | **0.240** | **0.435** | **0.497** | -0.018 | -0.038 | -0.003 | -0.027 | -0.034 | - |  |  |  |

|  |  |  |  |  |  |  |  |  |  |  |  |  |  |  |
| --- | --- | --- | --- | --- | --- | --- | --- | --- | --- | --- | --- | --- | --- | --- |
| PTB | **0.366** | **0.106** | **0.169** | **0.181** | **0.231** | **0.291** | 0.026 | 0.014 | -0.001 | 0.019 | -0.025 | 0.003 | - |  |
| SNQ | **0.352** | **0.090** | **0.153** | **0.165** | **0.211** | **0.273** | 0.022 | 0.009 | 0.002 | 0.016 | -0.027 | -0.004 | -0.049 | - |
| MLP | **0.532** | **0.086** | **0.141** | **0.201** | **0.377** | **0.446** | -0.016 | -0.035 | -0.005 | -0.030 | -0.030 | -0.048 | -0.005 | -0.008 |

**Table 3.** Hierarchical Analysis of Molecular Variance (AMOVA) on sequences of the mitochondrial control region for *Sphyrna lewini* in the Eastern Tropical Pacific. Based on Ɵ_ST_ pairwise comparisons. DF: degrees of freedom, SSD sum of squares, VC variance component, and % V percent of variance; two groups were defined: Northern Eastern Tropical Pacific (Mexico collection sites) and Central-southern Eastern Tropical Pacific (Guatemala, Costa Rica, Panamá and Colombia collection sites).

| | | DF | SSD | VC | %V | $\Theta$ | P value |
| --- | --- | --- | --- | --- | --- | --- | --- |
| Among regions | $\Theta_{CT}$ | 1 | 23.501 | 0.129 | 23.68 | 0.2367 | <0.0001 |
| Among populations within regions | $\Theta_{SC}$ | 13 | 6.351 | 0.002 | 0.49 | 0.0064 | 0.4175 |
| Within populations | $\Theta_{ST}$ | 420 | 173.675 | 0.413 | 75.84 | 0.2417 | <0.0001 |

### Microsatellite loci

A total of 169 individuals from three coastal sampling sites in Central America (GUA, COS and PAN), were genotyped at 14 microsatellite loci. MICRO-CHECKER provided evidence of null alleles on four loci (Sle18, Sle25, Sle53 and Sle77), which were duly removed from further analysis. Loci Sle13 and Sle27 were found to be linked (p < 0.05, Fisher’s method) after performing population-specific and global pairwise comparisons between loci to determine linkage disequilibrium, and the latter was also removed from further analyses. The remaining loci presented no significant deviation from Hardy-Weinberg equilibrium after Holm-Bonferroni correction and presented a total of 1.45% of missing data (sample with no interpretable pattern of DNA fragments after PCR amplification). The genetic relatedness estimates of Wang, showed the best performance with our data (r = 0.81), demonstrating an overall coefficient of r = −0.06. Relatedness tests performed using M-L relate found 22 pairs of FS (p = 0.01). Consequently, to avoid bias one individual of each pair was removed from further population structure analyses. Individuals sampled across all sites closely followed the distribution of values expected from unrelated pairs (Fig. 3). Unique alleles were found in loci Sle013, Sle033, Sle038, Sle054, Sle071, Sle081, Sle086, Sle089 and in all sampling sites (Table 1, S2).

**Figure 3.**
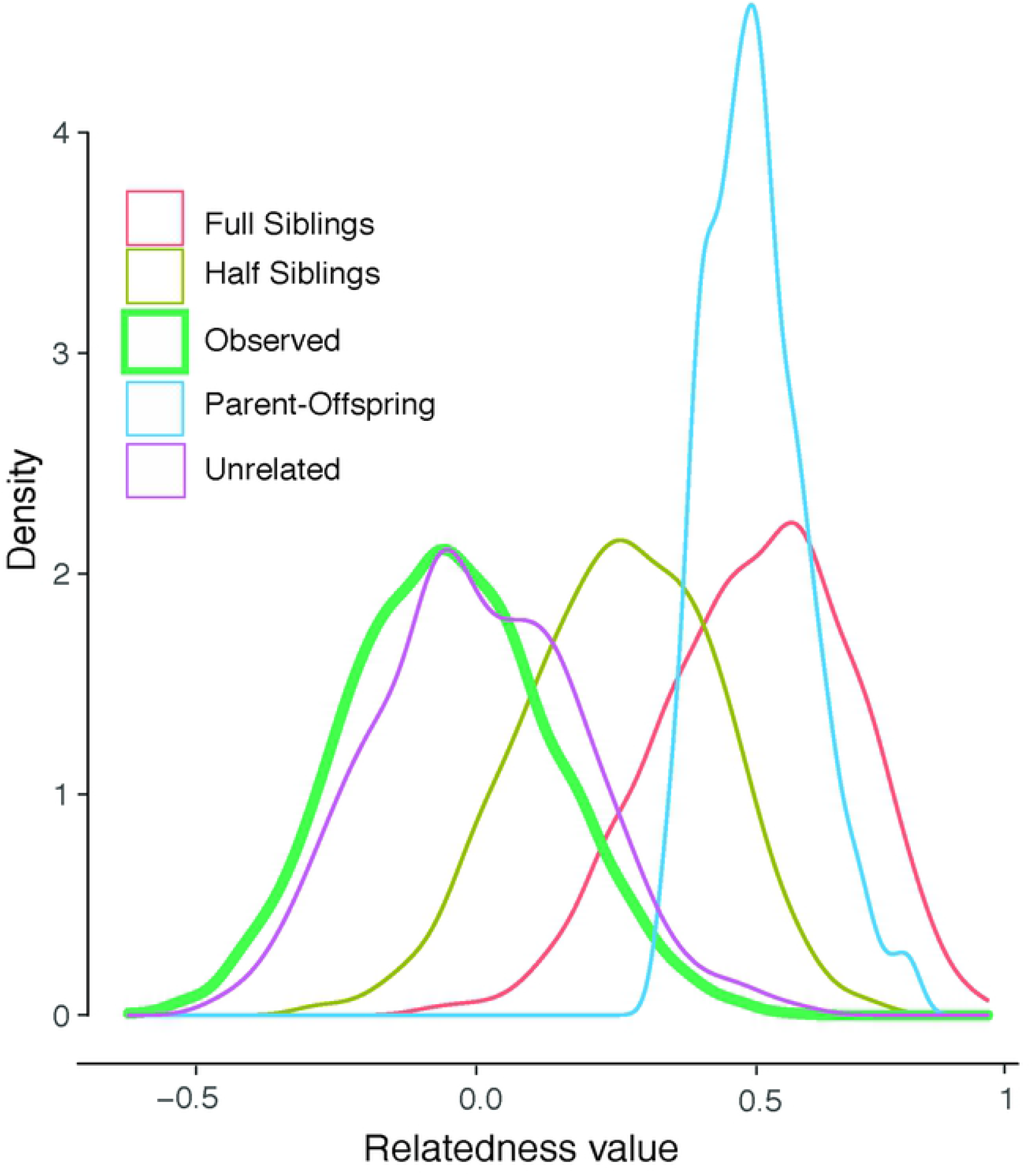
Distribution of pairwise genetic relatedness values for simulated pairs of individuals: Full siblings (FS), Half siblings (HS), Parent/Offspring (PO), and for observed pairs of individuals of *Sphyrna lewini* sampled in coastal areas of the ETP.

Genetic diversity metrics were similar between sampling sites (Table 1). The highest values of observed heterozygosity (Ho) and allelic richness (Ar) were found in Guatemala and the lowest in Panama (Table 1). Inbreeding coefficients (Fis) ranged from 0.02 to 0.10 (Table 1). Genetic diversity statistics were similar between the 9 loci analyzed (Table S2). Allelic richness across loci was 6.13 ± 4.06. The observed heterozygosity (Ho) ranged from 0.539 (Sle054) to 0.835 (Sle089). The inbreeding coefficients ranged from (Fis) 0.003 (Sle045) to 0.074 (Sle033).

Global genetic structure coefficients of D_EST_ and F_ST_ determined significant values of genetic differentiation for *S. lewini* at coastal sampling sites of the ETP (D_EST_ = 0.14, p < 0.05; F_ST_ = 0.054, p < 0.05). Pairwise comparisons of D_EST_ between coastal sampling sites were all significant and showed a greater differentiation between Guatemala and Panama (D_EST_= 0.222, p <0.05) and a lower differentiation for Costa Rica and Panama (D_EST_= 0.079, p < 0.05). F_ST_ values were concordant with D_EST_ values, showing less differentiation between Costa Rica and Panama (F_ST_ = 0.007) and greater differentiation between Guatemala and Panama (F_ST_ = 0.018). The DAPC conducted for the coastal sampling sites of the ETP show this same pattern of differentiation (Fig. 4A and Fig S1). Three groups were revealed by the STRUCTURE cluster analysis (K = 3) (Fig. 4B). Genetic distance was not correlated to the geographic distance between sites since the Mantel test revealed no significant IBD (p > 0.05.) Analysis of the extent and direction of gene flow showed no significant movement between coastal sampling sites. However, relative pairwise gene flow demonstrated higher connectivity between Costa Rica and Panama than between Panama and Guatemala (Fig 5). The genetic exchange obtained with this analysis coincides with the genetic population structure found in pairwise fixation indexes (D_EST_ and F_ST_) and the cluster analyses (STRUCTURE and DAPC) (Fig 4). Gene flow analysis including Cocos Island, demonstrated higher connectivity of this oceanic island with Costa Rica and Panama than with Guatemala (Fig S2).

**Figure 4.**
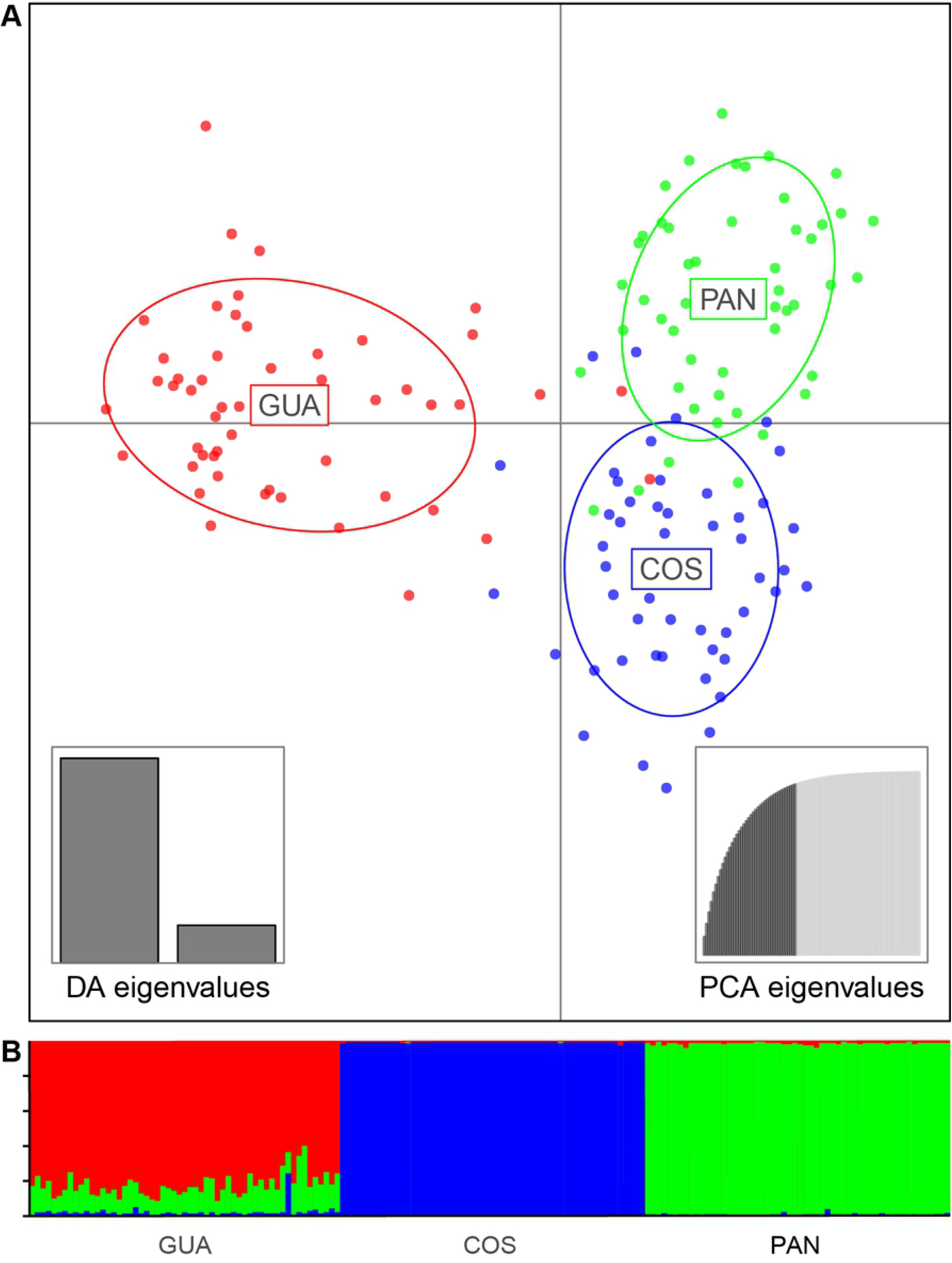
Population structure analyses from microsatellite genotypes of *Sphryna lewini* individuals in three sampling sites of the Eastern Tropical Pacific: Guatemala (GUA), Costa Rica (COS), Panama (PAN). **A)** DAPC plot from the first and second components of the nuclear microsatellite genotypes **B)** Genetic clusters inferred by STRUCTURE with K=3.

**Figure 5.**
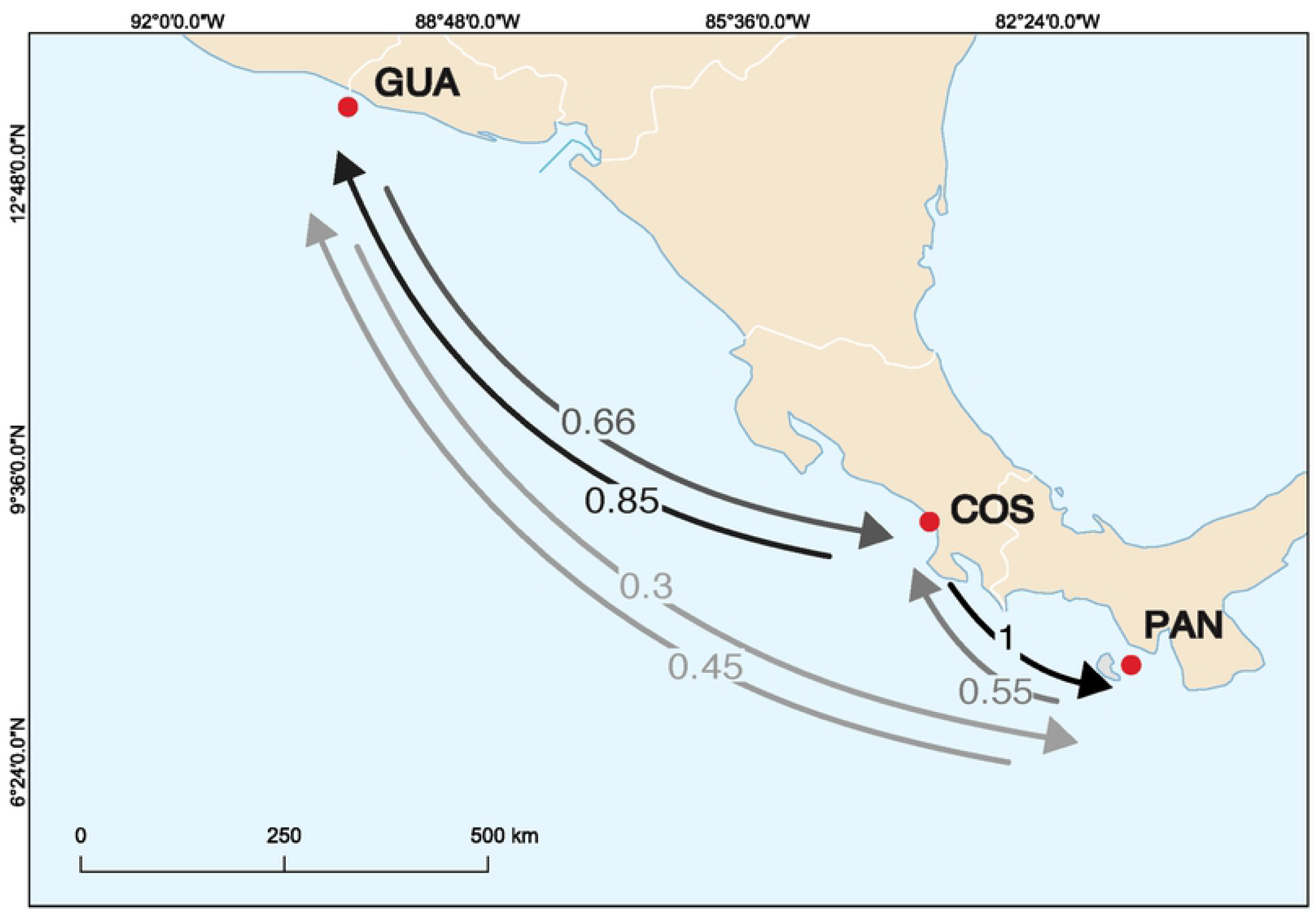
Contemporary gene flow estimated from 9 microsatellite loci genotypes with the divMigrate function. Arrows represent the relative number of migrants and estimated direction of gene flow between Guatemala (GUA), Costa Rica (COS) and Panama (PAN).

Overall observed average relatedness calculated from Wang (R= 0.0079) was significantly higher than would be expected by chance (R= −0.0391), indicating non-random relatedness in *S. lewini* individuals from each sampling site (Fig. 6). Additionally, the observed average relatedness in each sampling site was significantly higher than expected, indicating that individuals from these areas were more closely related within sampling sites than would be expected by chance (Fig. 6). No difference in average relatedness was observed between males and females (Fig. S3).

**Figure 6.**
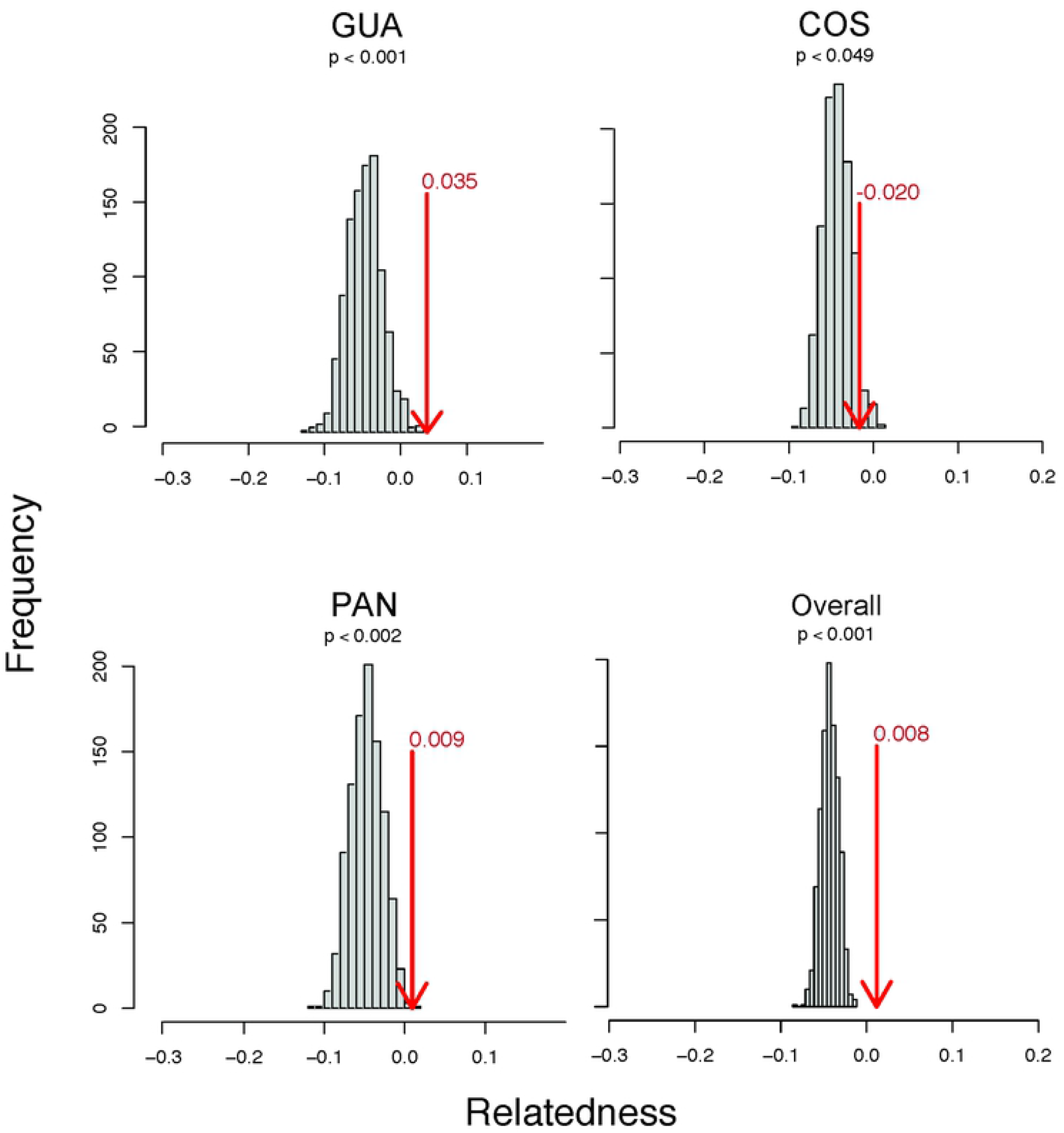
Expected distribution of average relatedness based on the Wang estimator of *Sphyrna lewini* in each sampling site and overall sampling sites using 1000 iterations. The average within sampling site relatedness examined within sampling site and overall sampling site is the statistic test (observed in a red arrow). The further away the statistic test is from the simulated bars, the greater the significance of the relatedness test.

## Discussion

Unravelling the genetic structure and connectivity is a fundamental step in the proposition and guidance of conservation and management strategies for threatened marine species. This is the first study to use a robust sample size to examine the fine-scale population genetic structure of this critically endangered shark species throughout the entire ETP. Patterns of genetic variation of the scalloped hammerhead shark *S. lewini* across multiple coastal areas within the ETP were assessed using both nuclear-encoded microsatellites and sequences of the maternally inherited mtCR.

### Genetic diversity

As with previous analyses of mtCR in the ETP (7,36,37), low levels of genetic diversity for *S. lewini* were found. This finding is consistent with the assumption that overexploitation of this species has led to the decline of its populations on a global scale (16). These levels of mitochondrial genetic diversity are comparable with those found in a recent study of this species in the Western Indian Ocean (hd= 0.232, n= 0.001) and Eastern Indian Ocean (hd= 0.362, n= 0.001) (66). Gene diversity, based on nucleotide and haplotype diversity, was highest in the Central-southern ETP, and lower in the Northern ETP. These differences coincide with the pattern observed among regions (Northern ETP and Central-southern ETP), which implies that the genetic signature of these regions may have resulted from an independent evolutionary history. In addition, nuclear microsatellite marker’s observed heterozygosity showed similar values to those previously reported for this species in the region Ho=0.703 (22), Ho= 0.770 (7). Observed heterozygosity in the ETP is similar to that reported for *S. lewini* in the Indian Ocean (Ho= 0.729) (22) and is higher than the heterozygosity values found in the Western Atlantic Ocean (Ho=0.580)(63). In the western North Atlantic Ocean, *S. lewini* suffered striking abundance declines, due to overexploitation (67,68), affecting the genetic diversity of this species in the region. When comparing the nuclear genetic diversity found in this study with that of other species, the values are similar to those reported for coastal sharks, including the bonnethead shark (*Sphyrna tiburo*) (Ho=0.59-0.69; (69)) and the blacknose shark (*Carcharhinus acronotus*) (Ho 0.66-68; (70)). In this study, a recent bottleneck effect was not detectable for *S. lewini*, most likely because only a few generations have passed since overfishing started. A non-detectable bottleneck effect was also reported for this species in the Western Atlantic Ocean (63).

### Population genetic structure

The mitochondrial DNA haplotype distribution of *S. lewini* revealed a pattern of differentiation between the Northern ETP and the Central-southern ETP. Genetic differentiation between these two groups is mainly due to an uneven distribution of the two common haplotypes that were found across all sampling sites. These results differ from the genetic homogeneity that has been previously observed for *S. lewini* in the ETP (7,9), which may be partially explained by the finer geographic sampling and larger sampling sizes used in this study. The entire Eastern Pacific is considered as a single, well-defined subpopulation of *S. lewini* (16,23), yet based on our results, this definition should be reconsidered. Additionally, the low level of mtDNA diversity observed suggests that the mtCR variation in *S. lewini* is insufficient to detect genetic heterogeneity at small scales. It is possible that using more mitochondrial markers or the complete mitogenome could provide a higher resolution, as demonstrated in the speartooth shark (*Glyphis glyphis*), the bull shark (*Carcharhinus leucas*) and the silky shark (*Charcharhinus falciformis*) (71–73).

The genetic break identified in our study is located between the boundaries of the Costa Rica Dome and the Tehuantepec bowl (74), suggesting that the seasonal dynamics of these systems generate oceanographic conditions that may have an impact on gene flow for *S. lewini* and other marine species. In the ETP, Rodriguez-Zarate et al. (2018) detected a similar pattern of genetic differentiation in the mtCR of the Olive Ridley sea turtle (*Lepidochelys olivacea*), a migratory marine species with similar life history traits as *S. lewini*, where Mexican nesting colonies were genetically differentiated from those in Central America. Their study determined the existence of two oceanographically dynamic but disconnected regions in the ETP, with a mixing zone located in southern Mexico (75). Pazmiño et al. (2018), also detected differentiation within the ETP region separating the galapagos shark (*Carcharhinus galapagensis*) mtCR sequences found in the Galapagos Islands from the mtCR sequences found in Mexico; this pattern is attributed to secondary barriers that have generated historical geographic isolation (76). A recent study on *S. tiburo*, a species that is closely realted to *S.lewini*, shows that magnetic map cues can elicit homeward orientation (77). This map-like use of the information of Earth’s magnetic field can offers a new explanation on how migratory routes and population structure of sharks can be maintained in marine environments.

The genetic homogeneity tests based on nuclear microsatellite loci revealed three genetically independent units: Guatemala, Costa Rica, and Panamá, with demonstrated limitations to gene flow between these coastal areas. A previous study conducted along the ETP coast, using the same microsatellite markers, found genetic differentiation in pairwise comparisons of areas with high sampling effort but not in areas with low sample sizes (7). Similar to our study, the authors found genetic structure between Tarcoles in Costa Rica (N= 40) and Santa Catalina (N= 46) in Panama (7). This pattern could not be observed between Tarcoles (N=40) and the Gulf of Panamá (N=9)(7), which highlights the importance of using robust sample sizes in genetic structure analyses. Despite the limitation of gene flow found between the three coastal areas, the greatest genetic similarity is observed between Costa Rica and Panama’s demes. Additionally, connectivity was detected between Cocos Island and the three coastal areas, and more gene flow is observed between this oceanic island and Costa Rica and Panama than with Guatemala. This is the first observation of genetic connectivity between Cocos Island and coastal areas of Central America, and is analogous to the gene flow found between the oceanic island of Malpelo and coastal areas of the Colombian Pacific (36). The observations of genetic differentiation between coastal nursery areas together with the genetic connectivity with oceanic aggregation areas of adults, suggest that *S.lewini* exhibits philopatry to fixed coastal areas in the ETP region. Adult females may undertake long-range migrations to oceanic islands within the ETP but return to specific parturition areas.

### Relatedness and natal philopatry

Inferring relatedness from genotypic data alone, remains a challenge and should be used with caution (78,79), nevertheless it provides insight into the potential mechanisms underlying fine-scale behavioral processes with long term consequences on population dynamics. Female fidelity to specific nurseries may define reproductive units if females are returning to the same location during each gestation cycle to give birth, leading to closer relatedness among juveniles from the same location than with individuals from surrounding areas (72). Individuals within nursery areas were found to be more closely related than expected by chance, thus suggesting that *S. lewini* may exhibit reproductive philopatric behavior within the ETP.

Given that *S. lewini* can undertake long-range migrations within the ETP (80), it can be inferred that the resulting population structure is not a consequence of limited dispersal ability. Moreover, all our sampling sites are potential nurseries for *S. lewini* in the ETP and the observed nuclear genetic structure does not support the relation of increased genetic differentiation with increasing geographic distance. This pattern has also been observed in the Atlantic Ocean, where the main factor driving population subdivision in *S. lewini* is reproductive philopatric behavior rather than oceanographic or geophysical barriers (Pinhal et al., 2020).

### Implications for conservation and management

These results offer new insights into the genetic diversity and connectivity of *S. lewini* in the ETP. Our fine-scale population genetic analysis revealed the existence of at least two subpopulations within the ETP, one in the Northern ETP and another one in the Central-southern ETP. The strong genetic partitioning found, urges the recognition of two different genetic units in the ETP; a region that was previously considered to be one distinct population segment of *S. lewini* (16,23). Additionally, coastal sites from Guatemala, Costa Rica and Panama were found to have different evolutionary dynamics, probably attributable to female philopatry.

The potential presence of philopatric behavior of *S. lewini* within the ETP emphasizes the need to develop more effective conservation approaches. All coastal sites along the ETP that could potentially serve as nursery areas for *S. lewini* are currently subject to illegal, unreported and unregulated fishing (36,81,82). Therefore, protection of these nursery areas is crucial for maintaining the genetic diversity, and consequently adaptive potential, of this critically endangered species (1). For a philopatric species, management measures that identify and protect parturition areas and unique localized genetic diversity could be crucial to avoid regional extinctions (34).

## Acknowledgements

We thank the National Council of Protected Areas of Guatemala for issuing the research license (license no. I-DRSO-001-2018), The Ministry of Environment of Panamá for the research permits (SEX/A-61-19 and SEX/A-108-17), the fishers from the community of Las Lisas Guatemala, Daniel Góngora and other fishers from the Punta Chame community for help in field work, Regina Domingo for sample collection in Punta Chame market, Alejandra Barahona director of the Center for International Programs and Sustainability Studies from Universidad Veritas for her support, members of Shark Defenders, Small scale fishers of Coyote and Bejuco, in Nandayure, Guanacaste, Costa Rica, The Alvaro Ugalde Scholarship issued by Osa Conservation, the National Secretary of Science and Technology of Panamá, and CONAGEBIO in Costa Rica for the research permits (CM-VERITAS-001-2021).

## Supporting information captions

**Table S1**. Localities, the total number (n) and accession number of mtCR gene sequences for *Sphyrna lewini* from the Eastern Tropical Pacific.

**Table S2**. Genetic diversity indexes of each microsatellite loci from *Sphyrna lewini* individuals in the Eastern Tropical Pacific. Ta: annealing temperature, Ho: observed heterozygosity, He: expected heterozygosity, Ar: allelic richness, Na: number of alleles, Ua: unique alleles, Fis: inbreeding coefficient.

**Table S3**. Geographic distribution and frequency of mitochondrial control region haplotypes of *Sphyrna lewini* individuals from the Eastern Tropical Pacific.

**Fig S1.** Densities of individuals in discriminant function 1 of 10 nuclear microsatellite loci genotypes of *S. lewini* in three collection areas of Eastern Tropical Pacific: Guatemala (GUA), Costa Rica (COS), Panama (PAN).

**Fig S2.** Contemporary gene flow estimated from 9 microsatellite loci genotypes with the divMigrate function. Arrows represent the relative number of migrants and estimated direction of gene flow between Guatemala (GUA), Costa Rica (COS), Panama (PAN), Cocos Island (ICO).

**Fig S3.** Distribution of the relatedness value of the Wang estimator in females and males of *Sphyrna lewini* overall collection sample sites.

**Fig S4.** Discriminant Analysis of Principal Components plot from the first and second components of nuclear microsatellite genotypes from *S. lewini* individuals in three sampling sites of the Eastern Tropical Pacific: Guatemala (GUA), Costa Rica (COS), Panama (PAN), Cocos Island (ICO).

